# Effects of Cr stress on bacterial community structure and diversity in rhizosphere soils of *Iris pseudacorus*

**DOI:** 10.1101/2022.08.11.503686

**Authors:** Zhao Wei, Zhu Sixi, Yang Xiuqing, Xia Guodong, Wang Baichun, Gu Baojing

## Abstract

Rhizosphere microorganisms play an important role in improving soil microenvironment, which contributes to plant growth under heavy metal stress. However, the effect of chromium (Cr) on plant rhizosphere bacterial community is still unknown. In this paper, sole-cultivated pattern, two-cultivated pattern and three-cultivated pattern, combined with 16S rRNA high-throughput sequencing technology, the effects of Cr stress on bacterial community structure and diversity in rhizosphere soil of *Iris Pseudacorus* were analyzed. The results showed that under Cr stress, *I. Pseudacorus* showed good tolerance and enrichment. However, under Cr stress, the Alpha diversity indices (Shannon, Chao and Sobs) of rhizosphere bacterial community decreased by 9.1%, 30.3% and 28.0% on average, respectively. The change of bacterial community was 22.6% due to Cr stress, and the common species of bacterial community decreased by 4.2%. Proteobacteria, Actinobacteria, Acidobacteria, Firmicutes and Gemmatimonadetes accounted for more than 78.2% of the total sequence. With the increase of plant diversity, Bacteroides and Pseudomonas appeared successively, and the abundance of the dominant species increased obviously. Through the symbiotic network diagram, it was found that the synergistic effect between dominant species in two-cultivated pattern was significantly enhanced, and the soil microenvironment was significantly improved. In conclusion, the results of this study will provide a reference for understanding the response of rhizosphere bacterial community to heavy metal Cr and the interaction between wetland plants and rhizosphere bacteria during wetland phytoremediation.

**Graphical Abstract:** 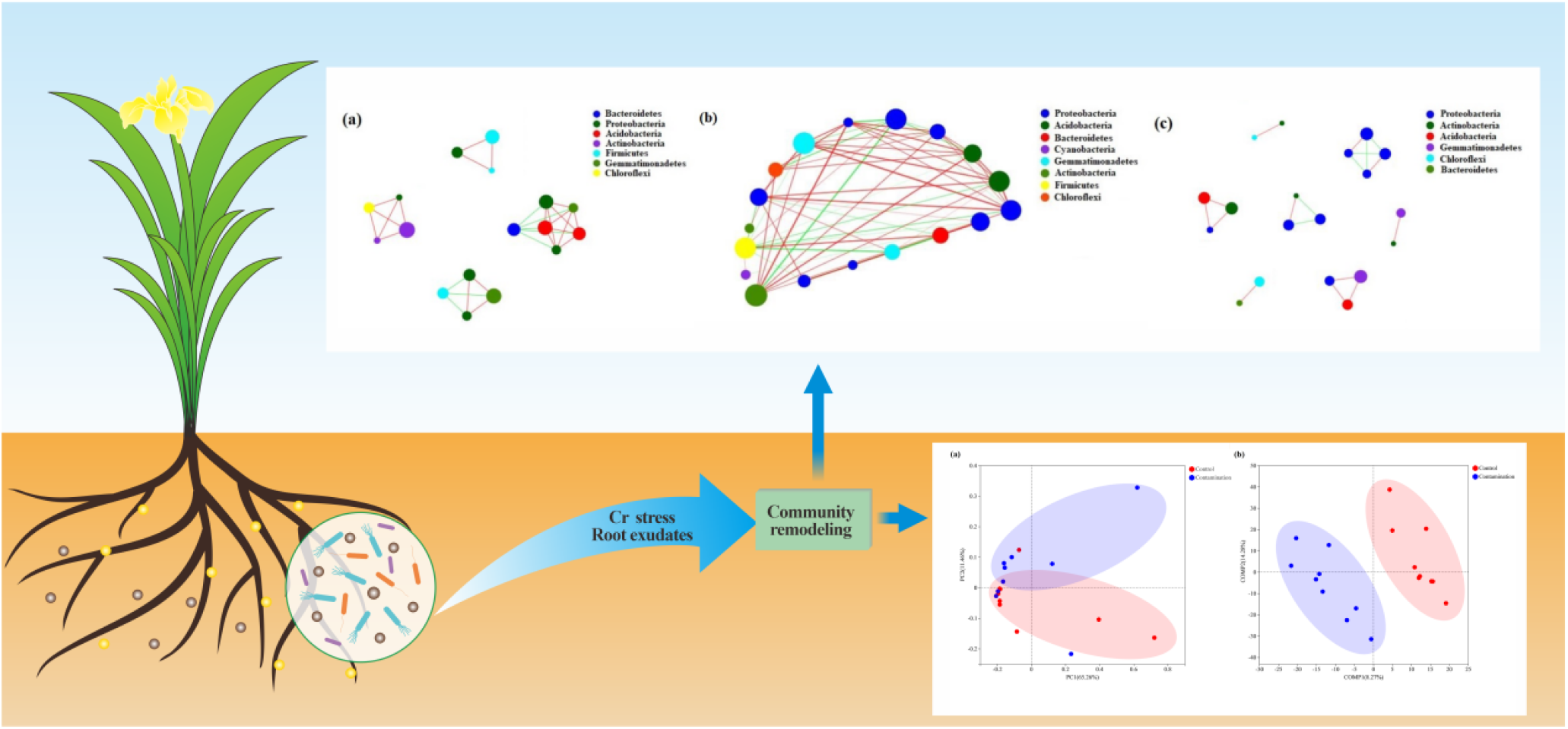

## 1. Induction

In recent years, due to the influence of human activities, heavy metal chromium (Cr) is widely distributed in soils and water bodies, and its environmental pollution has become a hotspot of concern (Xing et al., 2021). In addition to excessive exploitation of resources (Amit et al., 2018), Cr in the environment also mainly comes from metallurgy, electroplating, dyeing, tanning and other industrial fields (Zaheer et al., 2020). The most stable and common forms in nature are trivalent Cr(III) and hexavalent Cr(VI) (Kimbrough et al., 1999). Compared with Cr(VI), Cr(III) has weaker mobility, lower bioavailability and toxicity (Panda et al., 2005), and can also be used as an important component of glucose tolerance factor at low concentrations, which be beneficial to animal and human health (Losi et al., 1994). Due to its high fluidity and membrane permeability (Aashna et al., 2022), Cr(VI) has caused great harm to soil animals, microorganisms and plants (mainly manifested as soil degradation, microbial reduction and plant productivity decline (Peng et al., 2020). Ultimately, it will have a serious impact on human health through the food chain (Lin et al., 2020).

Cr hardly participates in any metabolic functions in plants, but it is potentially toxic and can produce toxic effects on plants (Wang et al., 2021). For example, as the Cr concentration gradient increases, Photosynthetic pigments (chlorophyll, carotenoids) (Rai et al., 2004), The synthesis of 1-aminocyclopropane-1-carboxylic acid and indoleacetic acid (Yaashikaa et al., 2022) will be inhibited, causes oxidative stress and enhances the synthesis of superoxide dependent on nicotinamide adenine dinucleotide phosohate (NADPH) (Guo et al., 2020). It affects the absorption of nutrient elements and water balance, and ultimately inhibits seed germination and root growth (Guo et al., 2020). The activity of antioxidant enzymes is reduced and membrane lipid peroxidation is enhanced, which leads to plant death (Hu et al., 2020).

The rhizosphere soil of plants is rich in microbial diversity. Rhizosphere microbiota is an important regulator of plant growth and stress resistance (Salas-Gonzalez et al., 2020). Rhizosphere microorganisms can increase carbon content in plant roots (Lange et al., 2015; Domeignoz-horta et al., 2020), nitrogen (N) (Zhang et al., 2019) and phosphorus (P) (Sundaresan et al., 2019) promote plant growth under adverse environment. It also affects the dynamic balance of plant antioxidant mechanism and adaptation to Cr stress (Guo et al., 2020). Moreover, rhizosphere growth-promoting bacteria (PGPR) can promote plant growth by dissolving mineral nutrients, secreting siderophore, indoleacetic acid and hydrocyanic acid, and release organic acids and biosurfactants to reduce the bioavailability and toxicity of Cr in soil, so as to improve plant tolerance. Promote growth on plants (Gupta et al., 2019). At present, Klebsiella variicola H12-CM-Fes@ (Yu et al., 2021a) and microbacterium Testaceum B-HS2 (Amina et al., 2019), Sporosarcina Saromensis W5 (Huang et al., 2021) and Bacillus sp. CRB-B1 (Tan et al., 2020) have been identified with high efficiency in removing chromium. Therefore, the use of bioremediation to reduce the toxic effect of Cr in the environment has a great prospect in the future (Muthusaravanan et al., 2018).

*Iris pseudacorus* is widely distributed in ponds and wetlands, and has been widely used in the remediation of Cr-contaminated soil due to its fast growth, high biomass, strong stress resistance, and strong tolerance and enrichment of heavy metals (Lin et al., 2018). Moreover, most wetland plants can fix Cr in their cell walls and vacuoles to reduce the toxicity of Cr to themselves (Wang et al., 2021). However, most studies focused on the mechanism of Cr removal by plants and the oxidative stress of Cr on plants, and there were few reports on the effect of Cr stress on the bacterial community structure and composition in the rhizosphere of *I. pseudacorus*. In this study, based on 16S rRNA high-throughput gene sequencing technology, we analyzed the physical and chemical properties of rhizosphere soil, the composition and diversity index of bacterial communities and the symbiotic network relationship between bacterial communities. And this study explored the mechanism of chromium stress on the structure and diversity of rhizosphere bacterial community. The results are of great significance for further understanding the response mechanism of rhizosphere bacteria to Cr stress in wetland plants and the mutual protection mechanism between *I. pseudacorus* and rhizosphere bacteria in Cr contaminated soil.

## 2. Materials and methods

### 2.1 Experimental design

*I. pseudacorus* were cultivated by greenhouse pot experiment with the flooded condition. Each pot contained 20 kg of soil collected from a karst mountain in southwestern China (106°37′36″E, 26°22′26″N), and multi-point mixed sampling was conducted to take soil samples from depths of 0-20 cm. Large chunks of weed stones were removed and passed through a 2 mm sieve. The pot of sole-cultivated pattern (IPI), two-cultivated pattern (IPII), three-cultivated pattern (IPIII) were designed as Control group (original soil samples without Cr contaminated) and Contaminated group (the exogenous addition of 0.1 mmol L^-1^ K_2_Cr_2_O_7_ solutions made Cr(VI) content to be 200 mg kg^-1^ in the soil), and the un-planted blank samples (CK), Each group had three replicates. Details of experimental grouping arrangement were shown in Fig. S1. The greenhouse ensured a constant temperature of 25 °C and moisture content of 50%, which reduced the interference of micro-meteorological factors on plant growth.

### 2.2 Sampling and chemical analysis

The pot culture started from the seedling (20-30 cm), and Cr(VI) was added to the pollution group after 30 days of domestication. After 3 months of pot experiment, destructive samples were taken. Randomly selected plants in potted plants were removed from the soil and collected the soil attached to the primary root (served as rhizosphere soil), and refrigerated to preserve the selected plants (Zhang et al., 2017); CK groups were potted to take soil samples of 0-5 cm from the upper layer. All of the samples were taken three times each time, totally 21 samples. Finally, samples were frozen at -20 °C for DNA extraction, high throughput sequencing and determination of Cr content.

The Cr contents of plants were measured by atomic absorption spectrophotometer (Perkin Elmer Analyst 800, USA). Soil physicochemical properties (SOM、pH、EC) were analyzed by the methods of Soil Agrochemical Analysis (Bao, 2000). Total genomic DNA from the rhizosphere soils were extracted and 16S rRNA sequenced following the manufacturer instructions and sent Majorbio Bio-pharm Technology, Shanghai for sequencing (Wang et al., 2021).

### 2.3 Statistical analysis

Normality and homogeneity of data were tested by KolmogorovSmirnov and Levene’s test. The data that did not obey a normal distribution were transformed by the natural logarithm. The weighted UniFrac distance algorithm was used to analyze the principal coordinates (PCoA), and the least square discriminant method (PLS-DA) was used to analyze and map (Wang et al., 2021). Analysis of variance (ANOVA) and the Mann-Whitney U test The significance of soil physicochemical parameters, plant physiological parameters and diversity index under chromium stress were tested. All statistical analyses were performed using SPSS statistical package (26.0) and Origin (2021). All the figures were prepared using Adobe Illustrator CC 2019 and Adobe Photoshop CC 2019 and Origin 2021.

## 3. Results

### 3.1 Soil physical and chemical properties and plant physiological parameters

In this study, after exogenous chromium addition, EC and pH values in soils increased by 16.6% and 5.0% respectively compared with the control group, among which, chromium treatment significantly increased SOM content in soil by 9.2% (Figure 1). In addition, the physiological parameters of plants were significantly changed after chromium was added. The Fresh weight of plants decreased by 45.1% on average, and the chromium content in roots and leaves increased by 97.1% and 99.5%, respectively (Figure. 1). It is noteworthy that after Cr stress, the Cr content in leaves of *I. pseudacorus* was significantly higher than that in roots, and the transfer factor reached 2.01. In conclusion, although Cr stress can destroy plant growth environment and inhibit plant growth, *I. pseudacorus* has a good enrichment effect and a certain tolerance to Cr.

**Fig. 1.**
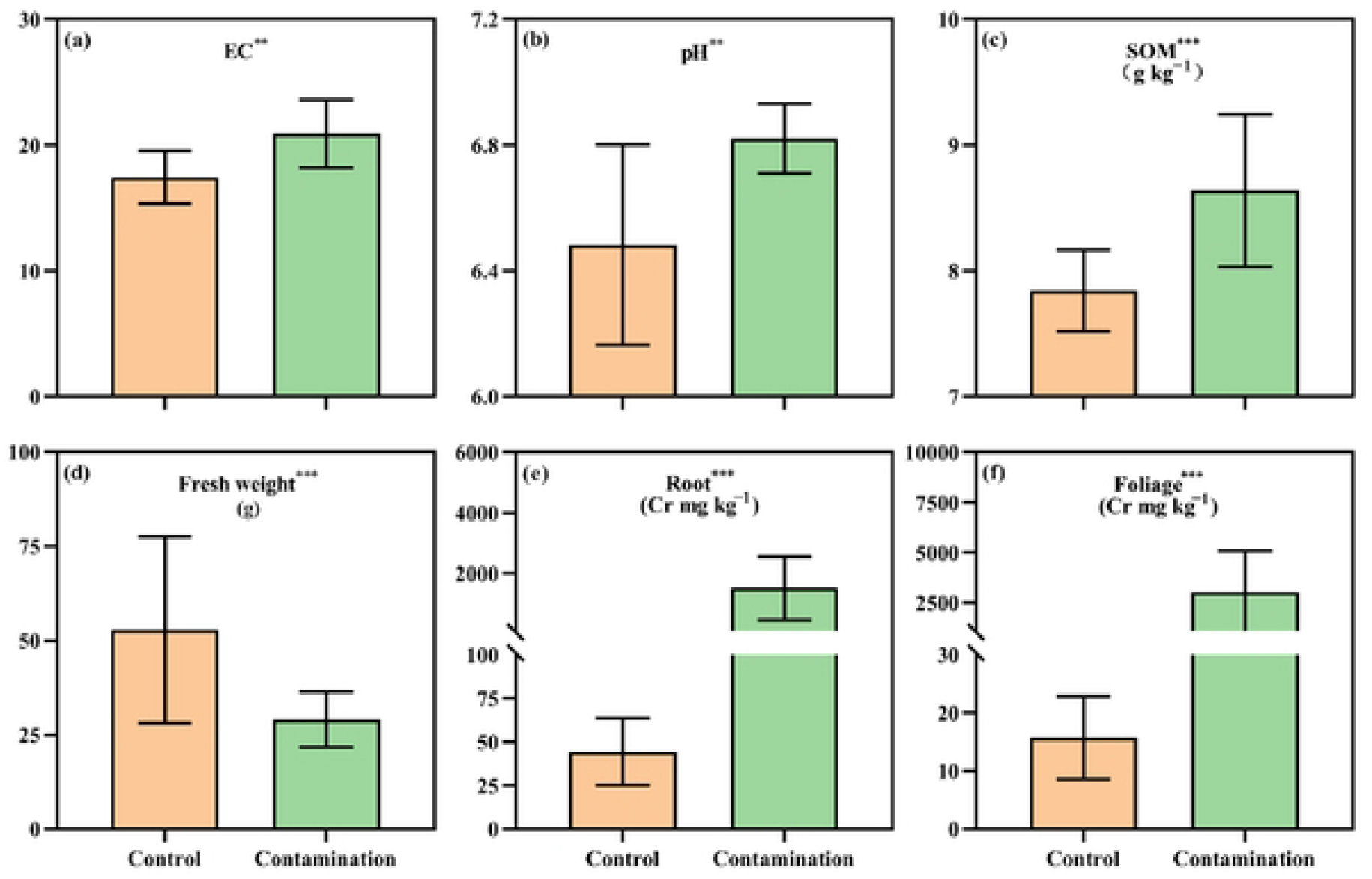
Plant physiological parameters and chemical properties of soil under Cr stress (a. EC; b. pH; c. SOM, d. Fresh weight; e. Root chromium content; f. Chromium content in foliage; *p < 0.05, **p < 0.01, and ***p< 0.001). The results showed that there were significant differences in physicochemical properties between the control group and the polluted group. Error bars refer to standard errors. Control contains treatments of IPI, IPII and IPIII, contamination contains treatments of CrIPI, CrIPII and CrIPIII.

### 3.2 Diversity index of soil bacterial community

In this study, unrecognized gene base sequences and chimeras were removed from the samples by high-throughput sequencing technology, and the Alpha diversity index of rhizosphere bacterial community of *I. pseudacorus* was analyzed (Figure 2). A total of 1239,259 valid sequences were obtained, and 2756 bacterial taxa (i.e. OTUs) were identified, among which the coverage of OTU sequences was above 98%. In addition, dilution curves were drawn according to Alpha diversity index, and it could be seen that the slopes of all sample curves were close to saturation (Figure. S2). Under Cr stress, the rhizosphere bacterial community Shannon index, Chao index and Sobs index of *I. pseudacorus* decreased by 9.1%, 30.3% and 28.0% on average compared with the control group (Figure 2), respectively. It can be seen that exogenous Cr supplementation significantly reduced the Alpha diversity index of bacterial community in rhizosphere soils of *I. pseudacorus*.

**Fig. 2.**
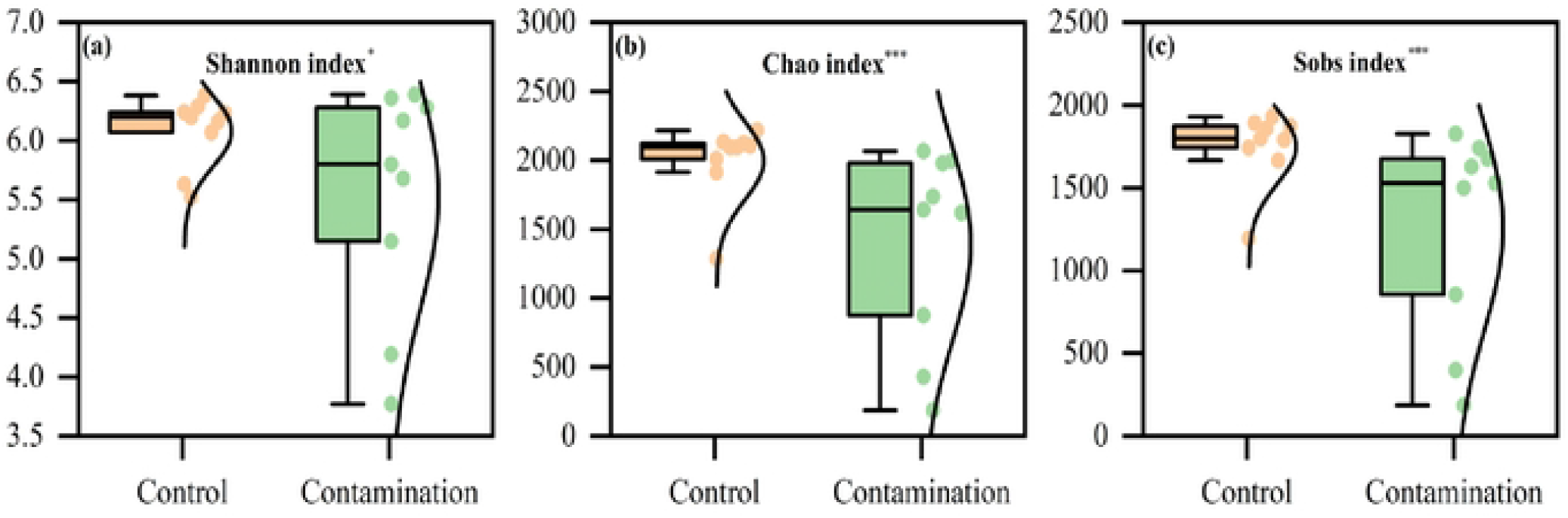
Comparison of bacterial community diversity and abundance index between control and contaminated groups (a. Shannon index; b. Chao index; c. Sobs index; *p < 0.05, **p < 0.01, and ***p< 0.001). The results showed that there were significant differences between the control group and the contaminated group. Error bars are outliers. Control contains treatments of IPI, IPII and IPIII, Contamination contains treatments of CrIPI, CrIPII and CrIPIII.

### 3.3 Soil bacterial community structure and composition

Veen diagram showed that the number of species co-existed in the control group and the polluted group was 17.0% and 16.4% respectively (Figure. S3a and b). Compared with the control group, the number of species co-existed in the Cr polluted group decreased by 4.2%. PCoA based on OTU taxonomy level showed that the bacterial community structure of the control group and the contaminated group were significantly different, and there were significant differences in spatial and temporal distribution pattern (Figure 3a). Among them, the difference explanation rates of the first principal component (PCoA1) and the second principal component (PCoA2) were 65.3% and 11.5%, respectively. Further PLS-DA analysis showed that the bacterial community changed by 22.6% due to chromium addition (Figure 3b). These verified that the addition of exogenous chromium could significantly affect the spatial pattern and composition of bacterial community.

**Fig. 3.**
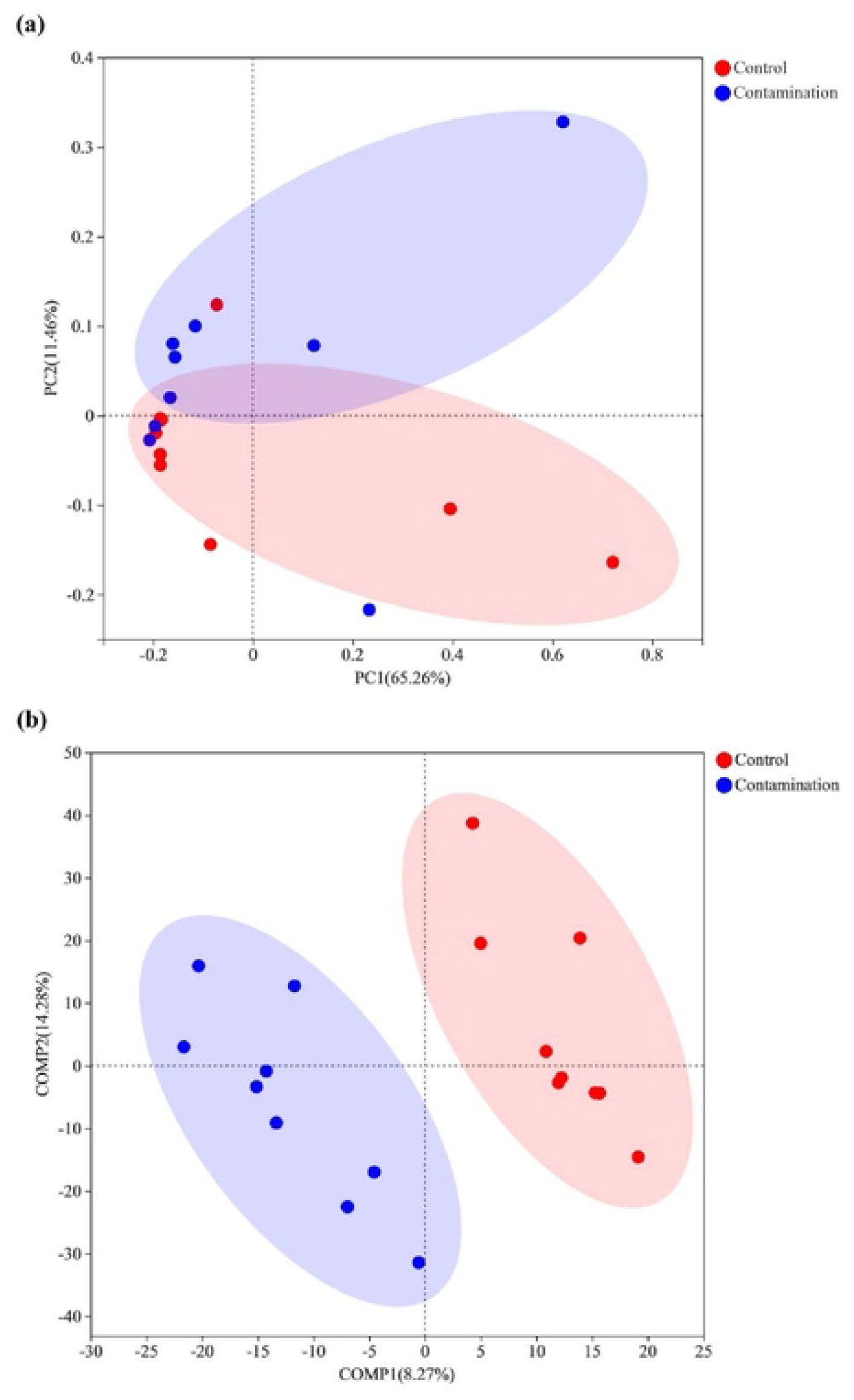
PCoA analysis on weighted UniFrac distance (a), PLS-DA analysis (b), which based on OTUs at a 97 % similarity level. PLS-DA performance of differentiation judges the potential interference factors, followed by OTUs at a 97% similarity degree. Comp1 and Comp2 respectively represent the suspected affecting factors for the deviation of microbial composition. Control contains treatments of IPl, IPII and IPIII, Contamination contains treatments of CrIPI, CrIPII and CrIPIII.

In this study, a total of 2756 bacterial taxa were identified, belonging to 37 phyla, 88 classes and 798 genera. The dominant phyla were Proteobacteria (average relative abundance 38.9%), Actinobacteria (14.2%), Acidobacteria (9.2%), Firmicutes (8.0%), Gemmatimonadetes (7.9%) and Chlo Roflexi (7.5%), Bacteroidetes (5.9%) and Bacteroidetes (Figure 4a). Compared with the blank group, Bacteroides and Pseudomonas were the dominant newly emerged bacteria (Figure 4b). Proteobacteria showed an obvious upward trend compared with the control group and the blank group under Cr stress in two-cultivated pattern. In addition, Proteobacteria, Actinobacteria, Firmicutes and Bacteroidetes showed an obvious upward trend in overall abundance compared with the blank group in the process of increasing plant diversity. Acidobacteria, Gemmatimonadetes and Chloroflexi showed a downward trend. It should be noted that Bacteroides increased first and then decreased during the increase of plant diversity, and the abundance of Bacteroides was the highest in sole-cultivated pattern under chromium stress. Among them, the new dominant bacteria in two-cultivated pattern that showed an obvious upward trend under Cr stress was Pseudomonas.

**Fig. 4.**
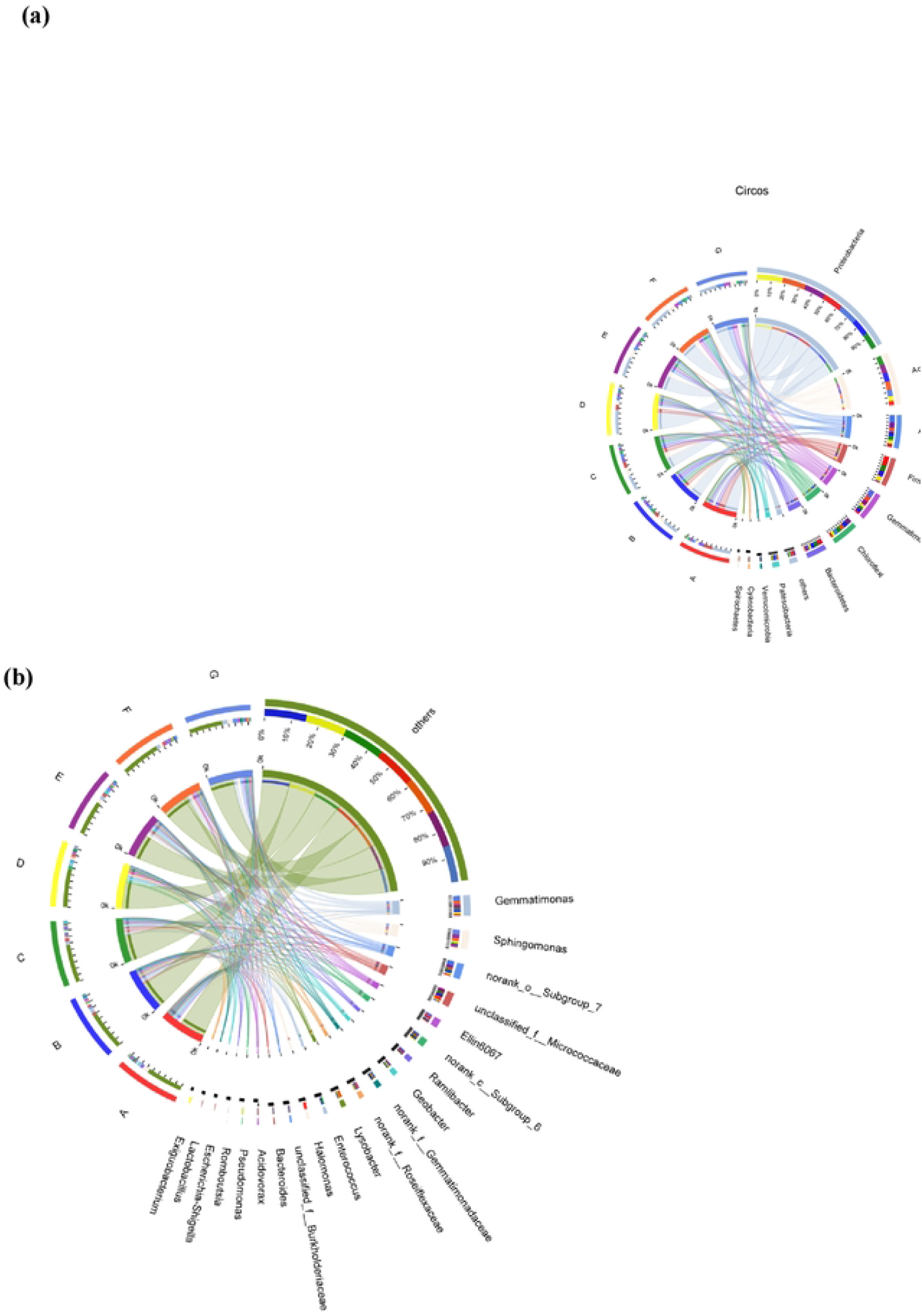
Structure and difference of bacterial community at class level under chromium stress(a:Phylum level; b: Genus level). The small semicircle (left half circle) represents the species composition in the sample. The color of the outer band represents the group from which the species are from. The color of the inner band represents the species, and the length represents the relative abundance of the species in the corresponding sample. The large semicircle (right half circle) represents the distribution ratio of species in different samples at the taxonomic level. The outer band represents species, the inner band color represents different groups, and the length represents the distribution ratio of the sample in a certain species.. A, C and E were the control group without Cr in *Iris pseudacorus* single, double and three plants, respectively. B, D and F were polluted groups with *I. pseudacorus* single, double and triple Cr treatments, respectively. G is for unplantcd and bulk soil (CK).

Further, the changes of dominant species under different cultivation modes were compared through the histogram of species abundance difference (Figure S4 a-d). We found that the abundance of Gammaproteobacteria, Actinobacteria and Alphaproteobacteria among the dominant strains of sole-cultivated pattern under chromium stress did not decrease, but showed an upward trend. Among them, Clostridia and Bacteroidia showed a decreasing trend. Under Cr stress in two-cultivated pattern, Gammaproteobacteria and Alphaproteobacteria showed the same upward trend, while Actinobacteria and Gemmatimonadetes showed a downward trend. Under Cr stress in three-Cultivated pattern, the abundance of Actinobacteria, Gammaproteobacteria and Gemmatimonadetes showed an upward trend, while Deltaproteobacteria showed a downward trend. On the whole, with the increase of plant diversity, the abundance of dominant species showed an obvious increase trend. Moreover, compared with the control group, the addition of plants significantly improved the abundance of dominant bacterial species in soil microenvironment.

### 3.4 Interspecific relationship of soil bacterial community

The symbiotic network diagram reflects the coexistence pattern of bacterial communities in specific habitats and explores the interspecific relationship of soil bacterial communities under Cr stress. Firstly, it can be seen from the colinear network diagram that the bacterial community structure among all samples is similar (Figure S5). Secondly, the symbiotic relationship between dominant species in sole-Cultivated pattern is weak, the competition is relatively prominent, and the symbiotic network is relatively scattered and not concentrated (Figure 5a). However, in the two-cultivated pattern, the symbiotic relationship between dominant species is significantly enhanced, while the competition relationship is significantly weakened. In addition, the symbiotic system formed among the dominant flora is closer and the symbiotic network is highly concentrated (Figure 5b). With the continuous increase of plant diversity, in the three-cultivated pattern, the symbiotic relationship between dominant species does not increase, but shows a downward trend. And the symbiotic network among species is more dispersed (Figure 5c). The results show that two-cultivated pattern is more conducive to improving soil microenvironment and creating a good living environment to resist Cr stress.

**Fig. 5.**
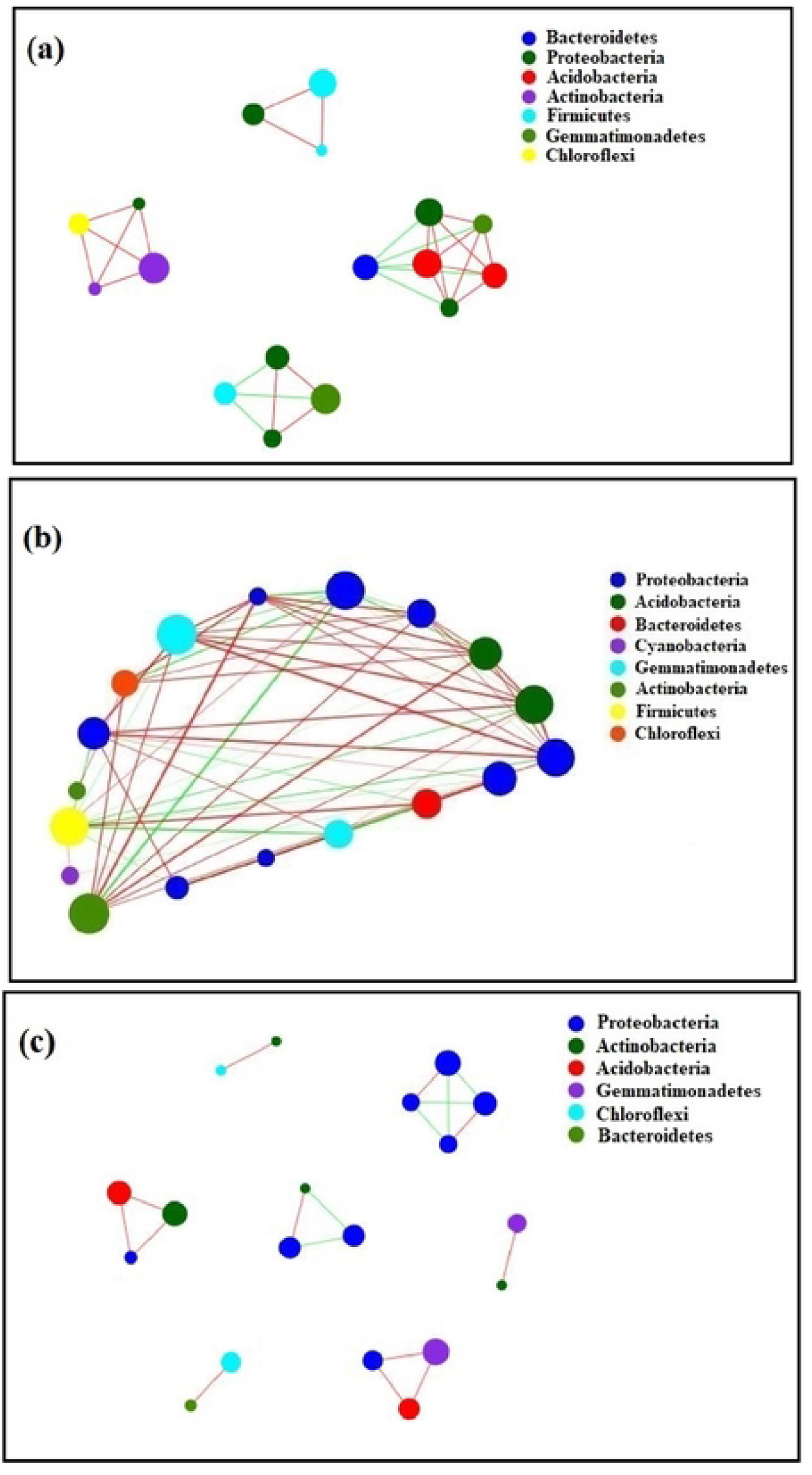
Symbiotic network diagram of soil rhizosphere microbial community (a: Sole plant model of *I. pseudacorus;* b: Double planting model of *I. pseudacorus;* c: Three planting model of *I. pseudacorus)*. The size of nodes in the figure indicates the abundance of species, and different colors indicate different species. The color of the line indicates positive and negative correlation, red indicates positive correlation, and green indicates negative correlation. The thickness of the line indicates the correlation coefficient. The thicker the line, the higher the correlation between species. The more lines, the more closely related the species is to other species.

**Figure 6.**
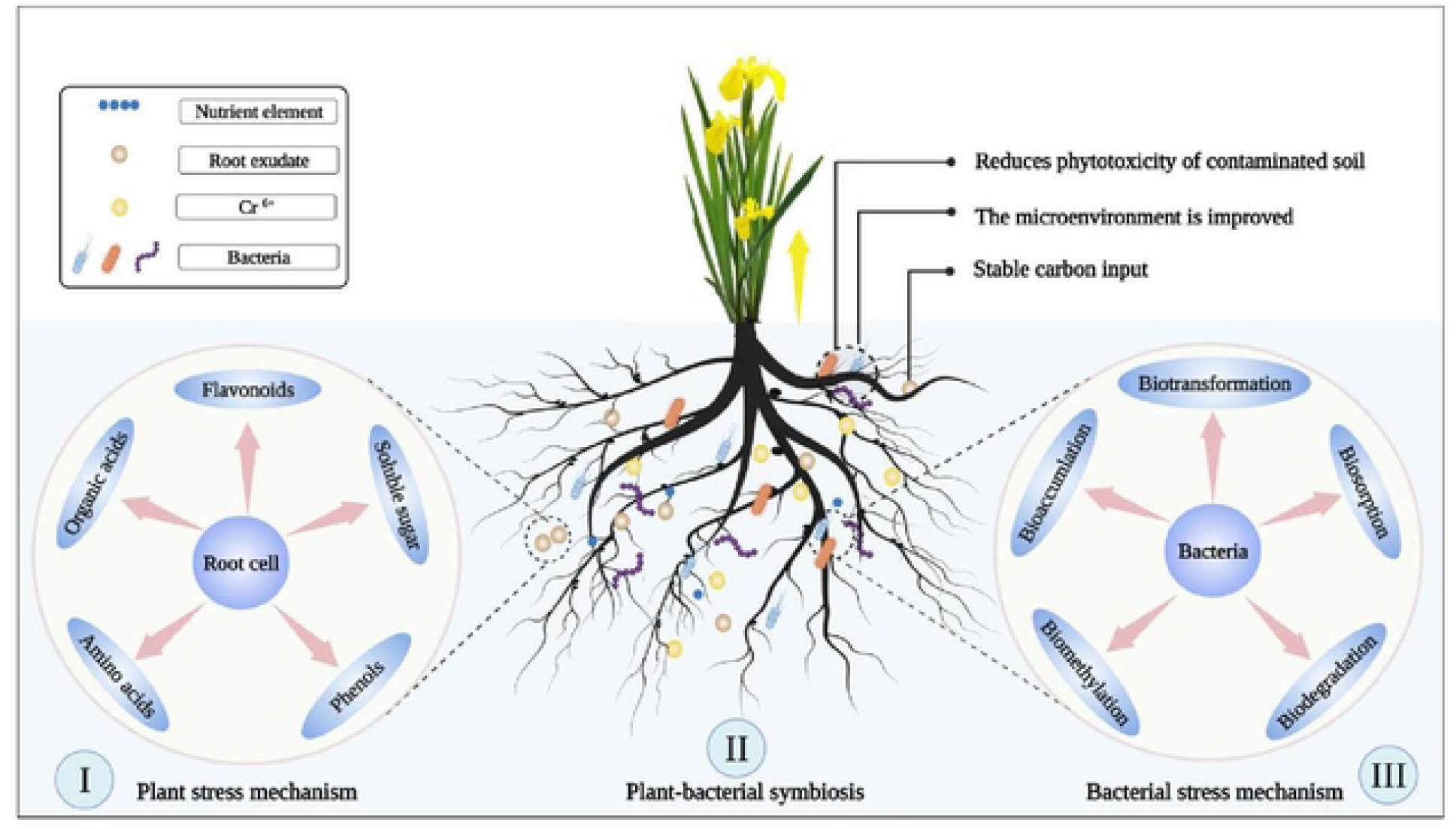
Schematic diagram of response mechanism of plant-microbe symbiosis system under Cr stress. (I) Plant stress mechanism; (II) Plant - bacterial symbiosis; (III) Bacterial stress mechanism. When treated with Cr, the plant will activate its own stress mechanism to respond to the external stress, thereby secreting a large amount of amino acids, organic acids, soluble sugar, and phenols. The content of organic matter in soil was increased, the microenvironment was improved, and the abundance of bacterial community was increased. When bacteria were subjected to Cr stress, It can reduce the toxicity of Cr by biotransformation, biosorption, bioaccumlation, biomethylation, biodegradation and other stress mechanisms, so as to reduce the toxicity of contaminated soil to plants.

## 4. Discussion

### 4.1 Responses of environmental factors of rhizosphere soils and *I. pseudacorus* to Cr stress

Excessive Cr in soil will inhibit plant growth and seriously threaten the health of soil ecosystem (Ao et al., 2022). Cr(VI) is characterized by high toxicity due to its strong bioavailability and mobility (Guo et al., 2020). It can inhibit plant photosynthesis, induce plant membrane lipid peroxidation and ROS production, thus affecting plant growth (Adhikari et al., 2020). In this study, compared with the control group, pH and EC values of rhizosphere soil physical and chemical properties showed an obvious upward trend with the addition of Cr(VI) (Figure 1), because soil properties (pH and EC) can significantly affect the adsorption effect of heavy metals (John et al., 2007). When the pH is low, the adsorption of heavy metals is reduced, resulting in the improvement of the bioavailability and mobility of heavy metals (Rieuwerts et al., 2006). Therefore, the increase of soil properties (pH and EC) may be related to the mechanism by which plants interact with their microenvironment to cope with heavy metal stress. Studies have proved that different pH conditions will significantly affect the reduction of Cr(VI) by microorganisms. When the optimum pH is reached, relevant functional genes will be significantly up-regulated.Then the pollutants can be completely removed, and the pathways for degradation and energy metabolism will be more abundant (Hua et al., 2020).

Due to the large amount of negative charge on the surface of organic matter, it plays an important role in the reduction of Cr(VI) (Praveen et al., 2019), so the presence of soil organic matter will significantly affect the properties of Cr. In this study, exogenous Cr significantly increased the SOM content (Figure 1). It may be that the plants under heavy metal Cr poisoning, can through the metabolism of the body itself produce chelating substances (such as organic acids, amino acids, protein, etc.). And these substances effectively alleviate the impact from Cr before entering into cells by the root secretion combining with heavy metals. At the same time, it can enhance the antioxidant defense mechanism in vivo and reduce the damage of reactive oxygen species produced under Cr stress (Zhong et al., 2019). It may also be related to the bacterial community in the soil, because the death of sensitive bacterial community under chromium stress can provide a large amount of organic matter to the soil (Sokol et al., 2022). And existing research shows that the SOM of the different functional groups, such as carbonyl and carboxyl, phenolic-OH etc. It can be combined with Cr(VI) (Shi et al., 2020), which was the main host phase for Cr (Saranya et al., 2021).

A large amount of Cr was accumulated in the roots and leaves of *I. pseudacorus* (Figure 1), which was due to the good tolerance and enrichment of heavy metal stress in wetland plants themselves, which could fix Cr in the cell wall, cell membrane and vacuole in the form of chelates or complexes to reduce its toxicity (Wang et al., 2021). However, under the high concentration of chromium stress, the stress mechanism of wetland plants was not enough to completely alleviate the oxidative stress caused by Cr, so the biomass of wetland plants decreased by 45.1% (Figure 1).

### 4.2 Effects of Cr stress on rhizosphere soil bacterial community structure and diversity

The structure and species diversity of rhizosphere bacterial communities play a very important role in soil ecosystem and plant regulation mechanism (Jin et al., 2018). In this study, we observed that the addition of exogenous Cr(VI) significantly decreased the Alpha diversity index (Sobs, Shannon, and Chao index) of the rhizosphere bacterial community in *I. pseudacorus* (Figure 2). Similar results have been observed by other researchers in studies of pinwheel rhizosphere microorganisms grown under Cr stress (Wang et al., 2021). This is because a large number of chrome-intolerant bacterial communities cannot alleviate the toxicity of Cr under the stress of high concentration of Cr, thus causing the damage of cell membrane function and their own metabolism, and dying due to the inability to maintain their own life activities (Sharma et al., 2021). It has also been observed that bacterial death may be due to competitive interactions between bacteria competing for resources in extreme environments (Sokol et al., 2022). When chromium enters cells through phosphate or sulfate carrier proteins, reducing enzymes and reducing substances in bacteria will reduce Cr(VI) to low-toxicity Cr(III), but during this period, reactive oxygen species and hydroxyl radicals will be produced in large quantities, thus inducing oxidative stress (Sharma et al., 2021). The strong oxidizing substances produced can cause DNA damage, which leads to cellular genetic variation and bacterial death (Guo et al., 2020), negatively affecting bacterial community diversity and abundance.

Cr stress significantly altered the spatial structure of bacterial communities in the rhizosphere soil of *I. pseudacorus* (Figure 3). Previous studies have shown that when the soil is contaminated with chromium, the bacterial community will exhibit obvious partitioning phenomenon, which significantly changes the spatial structure of the bacterial community in the soil (Yu et al., 2021b). This result has been confirmed in other studies (Wang et al., 2021; Wu et al., 2022 and Xiao et al., 2022). In this paper, PLS-DA (Figure 3) and Veen analyses (Figure. S3) proved that the addition of exogenous chromium could significantly change the structural composition and species number of bacterial communities. In conclusion, Cr stress can significantly reduce the diversity of rhizosphere bacterial community and change the structure of rhizosphere bacterial community, thus affecting the rhizosphere soils microenvironment.

### 4.3 Responses of rhizosphere soil bacterial communities to Cr stress and different cultivation patterns

In chrome-contaminated soils, previous studies have found the existence of Cr-reducing bacteria and tolerant bacteria, which play an important role in maintaining the stability of soil microenvironment and plant growth (Bhanse et al., 2022). In this study, it was found that Proteobacteria, Actinobacteria, Firmicutes and Bacteroidetes showed an increasing trend in the overall abundance compared with the blank group (Figure 4). With the increase of plant diversity, the abundance of dominant bacteria in soil microenvironment increased significantly (Figure 4). This may be related to the introduction of plants, because plants secrete organic substances (soluble sugars, phenolic compounds, flavonoids and organic acids) through roots during growth, which provide sufficient nutrients for the growth of rhizosphere bacterial communities and improve the microenvironment (Wang et al., 2021). The abundance of Proteobacteria, Gammaproteobacteria, Alphaproteobacteria, and Gemmatimonadetes increased after Cr stress (Figure S4). This is because Gemmatimonadetes and all Proteobacteria bacteria are gram negative bacteria. Their cell envelopes are composed of outer membrane (containing anion lipopolysaccharide, phospholipid and outer membrane protein) and peptidoglycan (He et al., 2020). Under the stimulation of Cr stress, it activates its own stress matrix and metabolic function, so as to release a large number of sugars and organic substances into the soil microenvironment to combine with heavy metals, reduce their bioavailability and toxicity, and create a good living environment. However, the abundance of Proteobacteria increased (Garavaglia et al., 2010). Actinobacteria, a typical gram-positive bacteria, plays an important role in the removal and reduction of Cr(VI), and has good tolerance to chromium, so that it can survive in chrome polluted soils (Ramesh et al., 2010).

In this paper, it was found that under the stimulation of chromium stress, two new dominant bacteria genera (Bacteroides and Pseudomonas) emerged (Figure 4). And Pseudomonas bacteria has a good reduction effect on Cr(VI) (Dogan et al., 2011). With the progress of technology, Pseudomonas sp. (Strain CPSB21) was obtained from chromium-contaminated wastewater by strain separation technology (Pratishtha et al., 2018), And it significantly promoted plant growth and antioxidant enzyme activity, and alleviated membrane lipid peroxidation. The results revealed that Pseudomonas NEWG-2 had a good effect on Cr(VI) removal by bioadsorption (Noura et al., 2020). Bacillus sp. CRB-B1 was an efficient Cr(VI) reducing bacterium (Tan et al., 2020).

The symbiosis network diagram in this paper showed that under sole-cultivated pattern and three-cultivated pattern, the synergism between dominant bacterial communities was weak, antagonism still existed, and the symbiosis network was relatively dispersed (Figure 5). This may be because under sole-cultivated pattern, plant diversity was less and carbon input to soil microbial community, which was not sufficient to construct a complete symbiosis system of rhizosphere microbial community (Lange et al., 2015). Under three-cultivated pattern, rhizosphere microbial community had stable carbon input, which may be because plants competed with each other to rob soil microbial resources, resulting in dispersed rhizosphere bacterial community and relatively dispersed symbiosis network constructed by bacterial community (Zhou et al., 2022). Among them, under two-cultivated pattern, not only the synergism between bacterial community was obviously strengthened, but also the symbiotic net formed was highly concentrated. It can be seen that under Cr stress, two-cultivated pattern was the optimal existence form (Figure 5).

## 5. Conclusions

The results showed that although there was a certain tolerance and enrichment of heavy metal Cr in *I. pseudacorus*, chromium still affected its environment and plant growth. In addition, the abundance and diversity of bacterial communities in the rhizosphere soils of *I. pseudacorus* were significantly decreased, and the spatial pattern of bacterial communities was significantly changed, affecting the composition of the rhizosphere bacterial communities. With the increase of plant diversity, the abundance of dominant species in soil microenvironment increased obviously, and beneficial bacteria appeared. Among them, two-cultivated pattern effectively changes the symbiotic relationship among the dominant species, significantly strengthens the synergistic effect between the dominant flora, and forms a more concentrated symbiotic network. The results of this study are helpful to further understand the effects of Cr stress on rhizosphere bacterial community, and provide reference for ecological remediation of Cr-contaminated soils.

## Acknowledgments

This study is financially supported by the National Natural Science Foundation of China (31560107), and by the Science and Technology Support Project of Guizhou province, China (Guizhou Branch Support [2018]2807).

